# Proper experimental design requires randomization/balancing of molecular ecology experiments

**DOI:** 10.1101/109280

**Authors:** Miklós Bálint, Orsolya Márton, Marlene Schatz, Rolf-Alexander Düring, Hans-Peter Grossart

## Abstract

Properly designed (randomized and/or balanced) experiments are standard in ecological research. Molecular methods are increasingly used in ecology, but studies generally do not report the detailed design of sample processing in the laboratory. This may strongly influence the interpretability of results if the laboratory procedures do not account for the confounding effects of unexpected laboratory events. We demonstrate this with a simple experiment where unexpected differences in laboratory processing of samples would have biased results if randomization in DNA extraction and PCR steps do not provide safeguards. We emphasize the need for proper experimental design and reporting of the laboratory phase of molecular ecology research to ensure the reliability and interpretability of results.

## Introduction

Ecological studies regularly ensure that the experimental setup is randomized and/or balanced. This allows to interprete results with respect to the original questions and to minimize the influence of confounding factors. The importance of randomized experimental setups (Fisher 1936) along with balanced designs (Student & Student 1938) is well-known. Consequently, such designs are enforced today in manipulative or observational ecological research (Hurlbert 1984).

This is often handled differently with laboratory experiments in molecular biology. By laboratory experiments we mean the laboratory processing (versus obtaining) of samples to generate quantitative molecular genetic data: DNA extractions, polymerase chain reactions, DNA sequencing, etc., in order to obtain haplotype frequencies, taxonomically informative marker gene counts, gene expression measures, SNP tables, etc. Early genome-wide association studies (GWAS) are examples of how basic experimental design may be ignored and what the consequences are: the analyses are expensive, but the obtained data cannot be interpreted (or are misinterpreted) due to confounding effects of laboratory procedures (e.g. Sebastiani *et al.* 2010). The early problems lead to the current recognition of randomized and/or balanced laboratory experimental designs in medical genomics (Yang *et al.* 2008; Leek *et al.* 2010; Lambert & Black 2012).

Complex and expensive molecular genetic datasets are increasingly generated in ecology. It is important that these data are generated appropriately since important conclusions and recommendations are drawn from them, often addressing issues of global importance for nature, society and economy. Randomization or balancing in laboratory experiments is essential to avoid batch effects and other nondemonic intrusions (see Hurlbert 1984). This issue has been already raised by Meirmans (2015) in a recent opinion paper on population genetics. Meirmans notes that “It is perfectly possible that such randomization is already practised in genotyping laboratories everywhere and I am simply unaware of it. […], if this is the case, this is nowhere evident in the literature”. We have similar impressions and the screening of one randomly selected 2016 issue of five relevant journals supports this assumption (Molecular Ecology, The ISME Journal, Ecology and Evolution, Journal of Biogeography, Soil Biology and Biochemistry, Appendix 1). Only two of the 59 relevant studies report some form of randomization during the laboratory processing of samples. This small literature survey is surely not representative of overall molecular ecology research, but the pattern is worrying since a simple Web of Science search for the keyword combination “molecul* AND ecol*” resulted in over 1740 hits only from 2016.

The omission of randomization in the lab may allow chance events to systematically influence results. Such chance events are common everytime and everywhere: electric fallouts happen, sudden flaws incapacitates lab personnel, DNA extraction kits are not delivered in time or have been stored inappropriate, just to mention some. If samples are processed in batches, the coincidence of these events confounds the results and renders interpretations unreliable. The potential diversity of such events is so high that nothing can protect against them except randomization of lab procedures, potentially in combination with balanced designs.

Hurlbert (1984) notes that most of the time chance events have immeasurably small effects on the results. However, by nature they are also completely unpredictable, both in frequency and effect size. Since molecular ecology studies mostly work with high observation numbers (thousands of SNPs over genomes, thousands of operational taxonomic units - OTUs - in hundreds of samples etc.), even small chance events may result in statistically significant results (Carver 1993). Here, we demonstrate this with taxonomically informative marker gene fragments amplified from environmental DNA (eDNA metabarcoding). The eDNA was preserved in lake sediments and provides a perspective on lake ecosystem history over several decades. We looked at three aspects of methodological or biological interest: extracted DNA concentration, PCR efficiency and community properties (Fig. 1). We evaluated several sources of variation: 1) expected laboratory biases (DNA extraction kit, Deiner *et al.* 2015; Barlow *et al.* 2016), 2) unexpected laboratory biases (this case a sudden change in lab personnel) and 3) an ecologically interesting predictor (either the age of the sediment or the effects of a power plant).

**Fig. 1.**
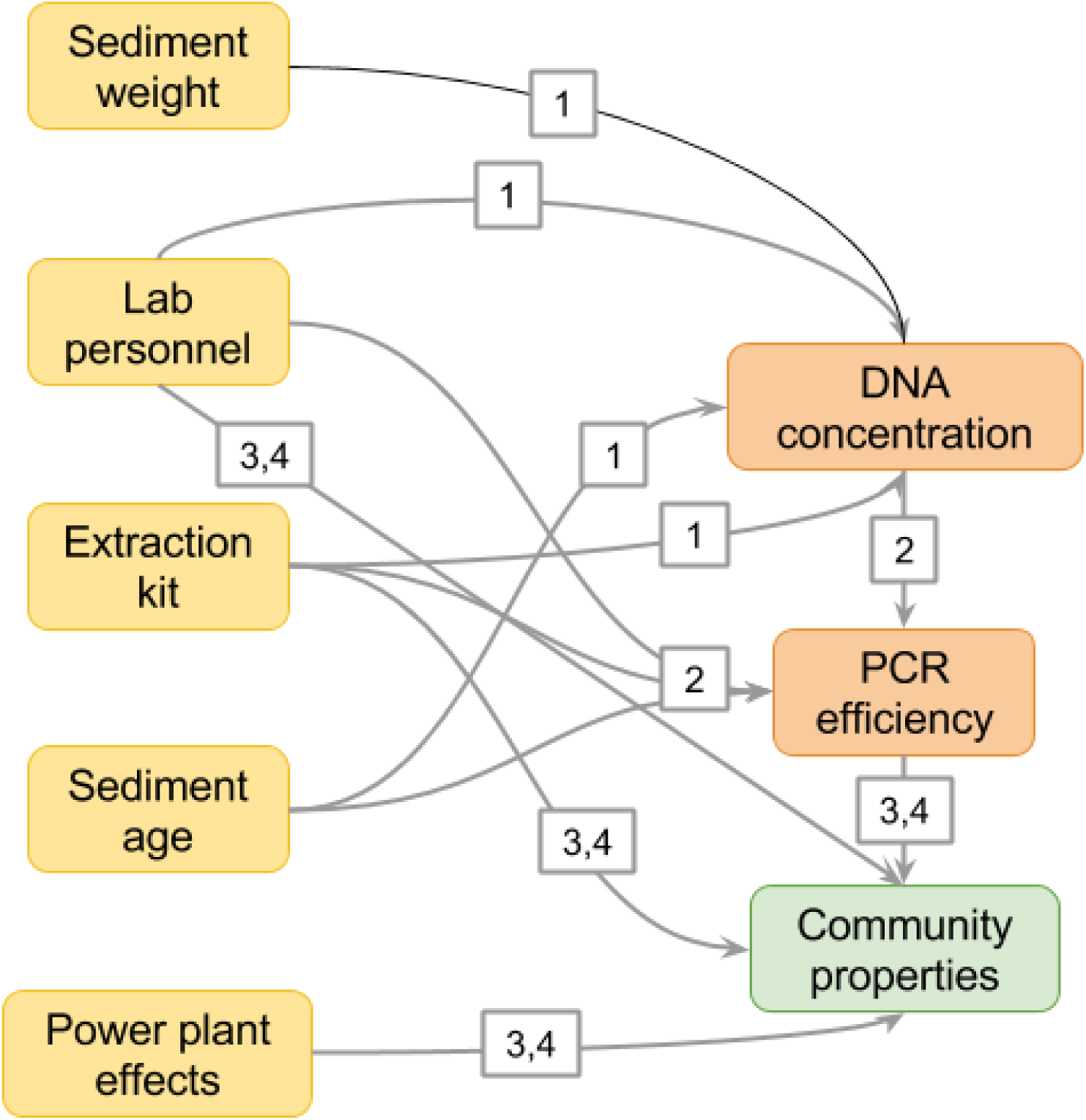
Analysis scheme with predictors of variation in high-throughput-sequenced eDNA data. Numbers on arrows refer to models of DNA concentration, PCR efficiency and community properties (see Materials and Methods). Yellow color marks variables that were used only as predictors in models. Orange variables were both predictors and responses in some of the models. Green marks variables that were only responses in models.

## Materials and Methods

### Sampling

Two sediment cores of the same location from Lake Stechlin were taken on May 14, 2015 with a gravity corer (UWITEC®, Mondsee, Austria) and Perspex tubes (inner diameter 9 cm, lengths 60 cm). Lake Stechlin (latitude 53°10‘N, longitude 13°02‘E) is a dimictic meso-oligotrophic lake (maximum depth 69.5 m; area 4.5 km^2^) in the lowlands of Northern Germany. GDR’s first nuclear power plant was built here between 1960-1966 and operated until 1990, connecting the lake with the nearby mesotrophic Lake Nehmitz and discharging its cooling water into Lake Stechlin. After coring, the cores were sliced immediately in the field in approximately 0.5 mm intervals. The first core was designated to eDNA. All sampling tools were H_2_O_2_-sterilized after cutting each horizon. Sediment for DNA extraction was taken only from the central part of the core to avoid contamination by contact with the corer’s wall. Samples were immediately stored in 15-ml Falcon tubes (NeoLab Migge GmbH, Heidelberg, Germany) at -20 ^o^C until DNA extraction. Horizons from the second core were used for organohalogenic pesticide measurements.

### Date approximation

Approximate dates were obtained by comparing DDT decomposition compound concentrations with sedimentation rates inferred with ^137^Cs (Casper 1994): 1.2 mm year^-1^ between 1986-1996 and 1.7 mm year^-1^ between 1963-1986. We assumed that DDT deposition started with World War II when a military training camp was operated near the lake and it effectively stopped in 1990 when agrochemical subventions of the GDR ceased with the reunification of Germany. The pesticide concentrations, sedimentation rates and inferred dates can be consulted in the file Stechlin_organohalogene.csv, deposited in Figshare (https://figshare.com/s/32dbca0a906c7f06449b, DOI: 10.6084/m9.figshare.4579681).

Halocarbon compound extraction was performed by shaking 300 mg freeze-dried sediment sample once in acetone and petroleum ether (40-60 °C) and then only in petroleum ether (40-60 °C), based on ISO 10382:2002. The clear supernatants were unified and vortexed after centrifugation, then a 10 ml aliquot was transferred to SPME amber screw top vials and evaporated under a gentle stream of nitrogen until dryness and dissolved again in 100 µL methanol and mixed with 10 mL of a 0.01 mol CaCl_2_ *2H_2_O / 3.4 mol NaCl salt solution. As internal standards 13C 2.4 DDT, 13C 4.4 DDT, α-HCH D6, Trifluralin D14, 4.4 BD D8, 4.4 DDE D8,13C HCB were used. Finally samples were extracted by Head Space Solid Phase Microextraction (SPME-HS) with a PDMS 100 fibre and analysed by GC/MS ion trap in selected ion monitoring mode (SIM). Separation and detection were accomplished using a Trace Ultra Gas Chromatograph (Thermo Fisher Scientific Inc., Schwerte, Germany) provided with a RTX-Dioxin 2 fused-silica capillary column with 0.25 µm film thickness, 0.25 mm ID and 60 m length coupled with an ion trap mass spectrometer in SIM (Thermo Fisher Scientific Inc., Schwerte, Germany).

### DNA extraction

We selected the youngest 21 horizons (the upper 13.5 cm of the core) for DNA extractions. Sample order was randomized before DNA extraction to minimize sampling biases. Four DNA extractions were carried out from each horizon with two commercial kits (two replicated extractions with both Macherey-Nagel NucleoSpin Soil - Macherey-Nagel, Düren, Germany, and MoBio PowerSoil - Carlsbad, CA, USA). The protocols of both kits were modified to specifically target extracellular DNA: instead of lysis, a saturated phosphate buffer was used to extract sediment-bound DNA (Taberlet *et al.* 2012). All four extraction replicates of a horizon horizon were performed in a row (see column extract_order in sample_infos.csv deposited in Figshare (https://figshare.com/s/32dbca0a906c7f06449b, DOI: 10.6084/m9.figshare.4579681). Four extraction controls (dH_2_O instead of sediment) were randomly included into the extractions. Extracted DNA concentrations were estimated on a Qubit 3.0 Fluorimeter (Thermo Fisher Scientific, Waltham, MA, USA).

### PCR amplifications

DNA templates were re-randomized before PCR setup. Four PCR negative controls (with dH_2_O instead of DNA template) and two positive controls (containing DNA from *Hypsiboas punctatus*, *Ponticola kessleri*, *Aspius aspius*, *Coregonus* sp., *Pacifastacus leniusculus*, *Aphanomyces astac*i, a parasitic Chytridiomycota, a saprotrophic Chytridiomycota, *Yamagishiella* sp., *Fragilaria crotonensis*, *Staurastrum planktonicum*, *Chaetomium* sp., *Lutra lutra*) were included. We used AmpliTaq MasterMix for the PCRs (Thermo Fisher Scientific, Waltham, MA, USA). We used general eukaryote primers that amplify a short fraction of the V7 region of the 18S gene region (Guardiola *et al.* 2015): forward - TYTGTCTGSTTRATTSCG, reverse - CACAGACCTGTTATTGC. The primers contained the Illumina sequencing primers (TCGTCGGCAGCGTCAGATGTGTATAAGAGACAG and GTCTCGTGGGCTCGGAGATGTGTATAAGAGACAG). The PCRs were run in 15 *μ*l reaction volume (AmpliTaq mastermix: 7.5 *μ*l, water: 4*μ*l, each 5 *μ*M primer 1*μ*l, DNA template 1.5*μ*l). The cycling conditions were 95 °C (10 min), 44 cycles of 95 °C (30 sec), 45 °C (30 sec), 72°C (30 sec), final extension at 72 °C (10 min). The PCR products were visualized on a 2% agarose gel and purified with Agencourt AMPure XP beads (Beckman Coulter GmbH, Krefeld, Germany).

### Multiplexing strategy and sequencing

We indexed all samples for multiplexed sequencing in a subsequent short PCR with primers that contained a fraction of the Illumina sequencing primer (TCGTCGGCAGCGTC and GTCTCGTGGGCTCGG), an eight-bp nucleotide index, and the Illumina plate adapters (P5: AATGATACGGCGACCACCGAGATCTACAC, P7: CAAGCAGAAGACGGCATACGAGAT). The final products are indexed, ready to sequence Illumina libraries. Index combinations and sequences are provided in the file multiplexing_indices.xlsx at Figshare (https://figshare.com/s/32dbca0a906c7f06449b, DOI: 10.6084/m9.figshare.4579681). The procedure follows the Illumina 16S metabarcoding protocol (Illumina 2016). This protocol eliminates index jumps during library preparation (although a few index jumps are still known to happen on the sequencing plate (Schnell *et al.* 2015). The indexing PCRs were run in 15 *μ*l reaction volume (AmpliTaq mastermix: 7.5 *μ*l, each 5 *μ*M primer 1*μ*l, PCR product 6.5*μ*l). The cycling conditions were 95 °C (10 min), 8 cycles of 95 °C (30 sec), 52 °C (30 sec), 72°C (30 sec), final extension at 72 °C (10 min). We checked the efficiency of each PCR run on a 2% agarose gel. The indexed libraries were purified with Agencourt AMPure XP beads (Beckman Coulter GmbH, Krefeld, Germany). The indexed libraries were mixed and purified on four QIAamp MinElute columns (Qiagen, Hilden, Germany). We did not normalize the PCR template concentrations to obtain a rough estimate of PCR and sequencing efficiency through the read numbers. Our sequencing kit potentially produces about 1 million paired-end reads with 2 × 150 bp length. Illumina sequencing was performed at the Berlin Center for Genomics in Biodiversity Research (www.begendiv.de) with the MiSeq sequencing kit v2 nano (300 cycles). Unprocessed sequence data were deposited in the European Nucleotide Archive as PRJEB19403.

### Sequence processing and data analysis

Raw sequence data were processed with OBITools (Boyer *et al.* 2015). Potential contamination and false detection biases were controlled for by following the recommendations of (Giguet-Covex *et al.* 2014; Boyer *et al.* 2015; Pansu *et al.* 2015) in R 3.3.1. (R Core Team 2016). All OBITools and R commands are documented in the file stechlin_analyses.pdf at Figshare (https://figshare.com/s/32dbca0a906c7f06449b, DOI: 10.6084/m9.figshare.4579681), with the full code accessible through the GitHub repository https://github.com/MikiBalint/LaboratoryDesign.git. Commands were run with GNU ‘parallel’ when possible (Tange 2011). The resulting OTU abundance table is provided in the stechlin_assigned_190915.tab file through Figshare (https://figshare.com/s/32dbca0a906c7f06449b, DOI: 10.6084/m9.figshare.4579681).

We fitted linear mixed-effect models with lme4 (Bates *et al.* 2015) on extracted DNA concentration, PCR efficiency and measures of diversity (the first three integers from Hill’s diversity series (Hill 1973)) to estimate the effects of potential laboratory biases and biological factors of interests. The first three Hill numbers correspond to species richness (H1), the exponent of Shannon diversity (H2), and the inverse of the Simpson diversity (H3). The identity of the sediment horizon was used as the random effect in these models. We used multispecies generalized linear models (GLMs) with the ‘mvabund’ R package (Wang *et al.* 2012) to investigate the effects of the predictors on community composition. The multispecies GLM cannot handle random effects. The community composition effects were visualized with a latent variable model-based ordination performed with the boral R package (Hui 2016). Both compositional analyses assume a negative binomial distribution of the data, accounting for the sparse and overdispersed nature of read counts (Bálint *et al.* 2016). The input data matrices are accessible through Figshare (https://figshare.com/s/32dbca0a906c7f06449b, DOI: 10.6084/m9.figshare.4579681).

The models can be written up as

1. conc ~ weight + kit + person + age + I(age^2) + (1|depth.nominal)
2. PCR efficiency ~ conc + kit + person + age + (1|depth.nominal)
3. diversities ~ PCR efficiency + person + kit + nuclear + (1|depth.nominal)
4. Community composition ~ reads + kit + person + nuclear,

where *conc* is the extracted DNA concentration, *weight* is the sediment weight used for DNA extraction, *kit* is the DNA extraction kit, *person* is the lab personnel, *depth.nominal* is the identity of the sediment horizon, *PCR efficiency* is estimated from HTS read numbers, *nuclear* is the operational period of the nuclear power plant (Fig. 1).

## Results

The results are summarized in Table 1 and Fig. 2. Regarding DNA concentrations, the DNA extraction kit (equivalent to the expected laboratory biases) accounted for most variation, followed by the age of the sediment horizon (biological signal) and the lab personnel (equivalent to the unexpected laboratory bias). The starting weight (amount) of the sediment had limited effects on the extraction efficiency and the effect of the lab personnel was marginally significant. PCR efficiency (evaluated as non-normalized HTS read numbers from PCR amplification) was mostly explained by the personnel identity (unexpected lab bias), followed by the DNA extraction kit (expected laboratory biases), the age of the sediment horizon, and the DNA template concentration used for the PCR. Here, the effect of the lab personnel was statistically significant. The most important contributors to variation in the first three Hill numbers consisted of PCR efficiency and the effects of the nuclear power plant. The DNA extraction kit contributed relatively little to the observed variation in the diversity indices. The nuclear power plant effects, however, represented the largest contributors to the explained variation in community composition, followed by the identity of the lab personnel and PCR efficiency. The DNA extraction kits contributed the least to the explained compositional variation (Fig. 3). The effects of the lab personnel were statistically marginally significant. Additional results and effect plots are available in file stechlin_analyses.pdf at Figshare.

**Table 1.** Summary of predictor contributions to variation. * statistically marginally significant result (p<0.1), ** statistically significant result (p<0.05), ^+^ statistical significance not tested.

|  | DNA<br>concentration | PCR<br>efficiency | H1 | H2 | H3 | Community<br>composition |
| --- | --- | --- | --- | --- | --- | --- |
| Sediment weight | 0.1<sup>+</sup> |  | - | - | - | - |
| Extraction kit | 12.9<sup>+</sup> | 2719604<sup>+</sup> | 672.6<sup>+</sup> | 197.9<sup>+</sup> | 67.8<sup>+</sup> | 906 |
| Lab personnel | 1.7<sup>**</sup> | 81413118<sup>**</sup> | 37.9 | 0.9 | 1.9 | 2018<sup>*</sup> |
| DNA concentration | - | 58<br>222<sup>+</sup> | - | - | - | - |
| PCR efficiency | - | - | 14834.7<sup>+</sup> | 156.3<sup>+</sup> | 72.3<sup>+</sup> | 1163<sup>**</sup> |
| Age/power plant effect | 1.8<sup>+</sup> | 198964<sup>+</sup> | 1758.3<sup>+</sup> | 867<sup>+</sup> | 435.5<sup>+</sup> | 5488<sup>*</sup> |

**Fig. 2.**
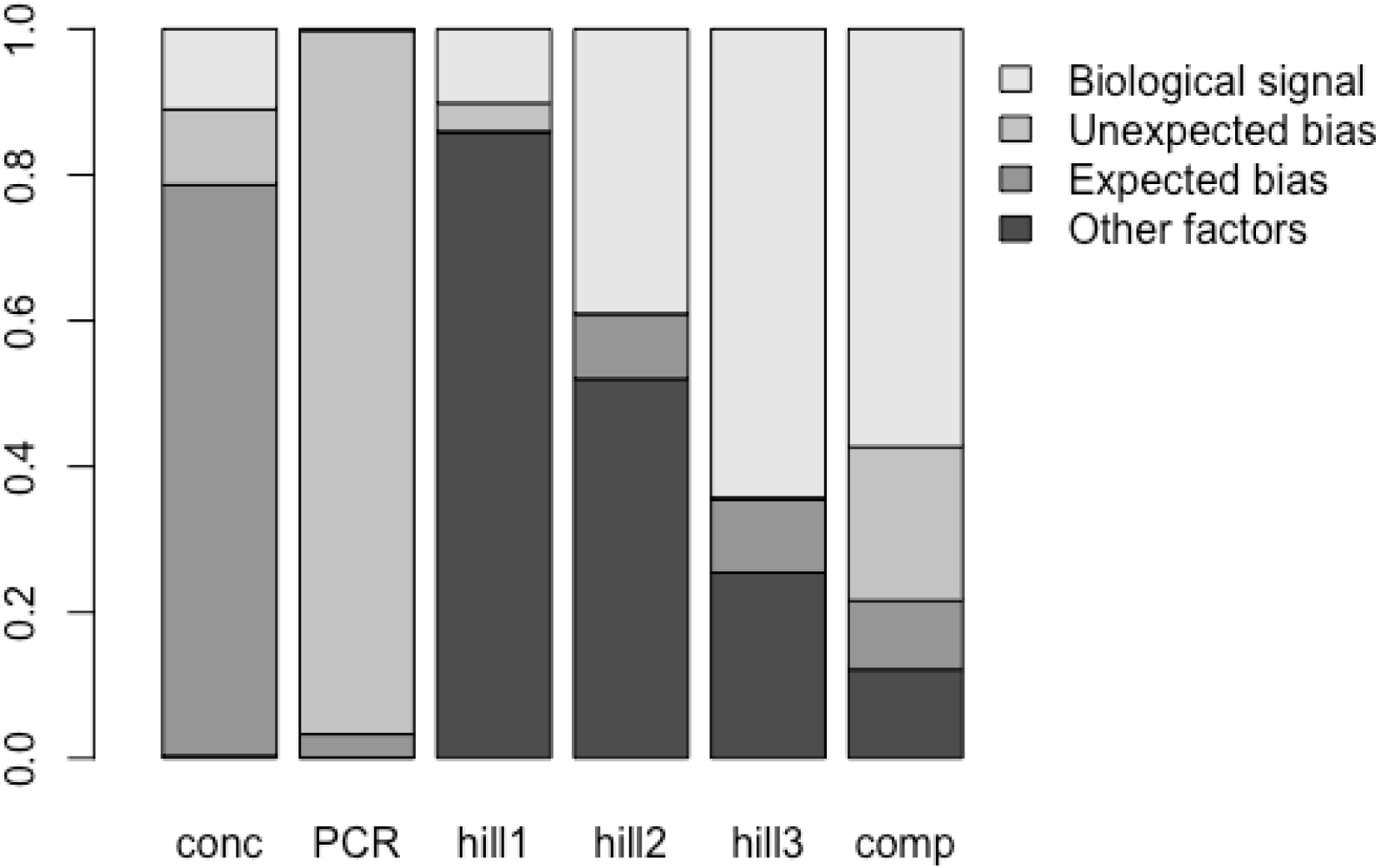
Partitioning of variance explained by expected and unexpected laboratory biases, and biological signal. The bars represent explained variance in DNA concentration (conc), PCR efficiency (PCR), diversity indices (hill1-3) and community composition (comp). Predictors: biological signal: effects of sediment age (conc, PCR) or the power plant operation periods (hill1-3, comp); unexpected bias: effects of laboratory personnel; expected bias: effects of DNA extraction kit; other factors: sediment weight (conc), DNA concentration (PCR); PCR efficiency (hill1-3, comp).

**Fig. 3.**
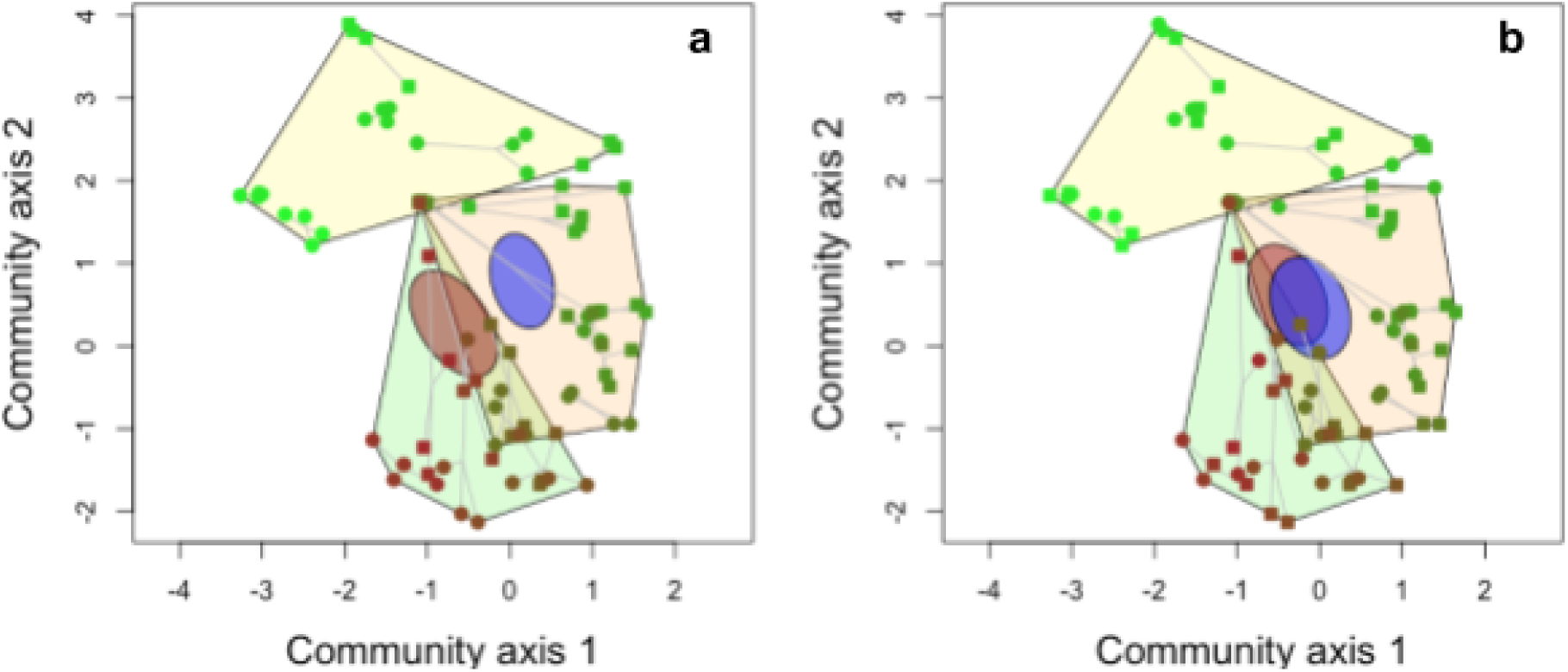
Compositional changes in historic communities explained by expected and unexpected laboratory biases, and biological signal. Points represent communities reconstructed from replicated DNA extractions from 21 sediment horizons, representing the last ~70 years of the lake’s history. Symbol color indicates age: dark brown are the oldest, light green are the youngest communities. Replicated DNA extracts of a horizon are connected by grey lines. The operational phases of the nuclear power plant are marked with hulls: green - before building the plant, orange - during power plant operation, yellow - after operation. a) symbols mark the effects of lab personnel on community composition and the two ellipses show the 95% confidence interval of the corresponding group centroids. b) symbols and ellipses mark the effects of the DNA extraction kits.

## Discussion

Our results demonstrate that nondemonic intrusions (Hurlbert 1984) in the laboratory may produce in strong, statistically significant effects that may severely confound results. Such effects render equivocal interpretations impossible if they coincide with effects targeted by the study. For example, interpretation of power plant effects on community composition would be difficult if samples are processed in batches and the sudden change in laboratory personnel coincides with a shift between operation periods.

We don't state that biases with comparable extent always appear in unrandomized, not balanced laboratory experiments, but they certainly have the potential to do so. This is clear in our example: the effects of unexpected laboratory biases exceed the effects of known lab biases (DNA extraction kit effects) and biological signal in several models (Fig. 2). Such effects potentially influence all molecular ecology studies and threaten the interpretability of results. Their importance and extent is widely known in biomedicine (Yang *et al.* 2008; Leek *et al.* 2010; Lambert & Black 2012) and needs to be urgently considered in molecular ecology.

Generally, randomization of samples before major laboratory steps (extraction, PCR, sequencing) is simple and low-cost. The only case where this might be disputable is the processing of highly contamination-prone materials where it is almost a lab rule that DNA extraction is performed consecutively from the most contamination-prone toward the least contamination-prone samples (although to our knowledge the validity of this still needs to be tested). Obviously, nondemonic intrusions (including contamination) in the laboratory easily become collinear with the processing order and this makes biological signals difficult to interpret (Salter *et al.* 2014).

We recommend the followings: first, researchers involved in molecular ecology labwork need to properly design and report laboratory procedures. Guidelines in biomedicine exist and may be readily adapted (Masca *et al.* 2015). Second, ecologists who rely on molecular data generated by laboratory personnel or companies must ensure (and should not take for granted) that principles of experimental design are followed in the laboratory. This is the easiest when giving samples to a lab since the ecologist can already rearrange and relabel his/her samples (but controls of PCR, sequencing, orders, etc. may require further communication). Third, editors and reviewers of manuscripts and grants should enforce the reporting of laboratory experimental design. This is as much necessary for reproducible research as the proper presentation of sampling schemes, details of manipulative experiments and data analysis. We do not intend to provide a list of important laboratory biases since there are potentially infinite variations. Therefore, molecular ecologists must ensure randomization or properly balanced designs in every step of laboratory work and present the details. There is no excuse for avoiding this since more and more globally important decisions require reliable molecular ecology data in nature and biodiversity conservation.

## Acknowledgements

The authors express gratitude to the working community of the Senckenberg Conservation Genetics Group (Gelnhausen, Germany) for hosting and supporting the labwork. Claudia Wittwer, Silke Van den Wyngaert, Martin Jansen, and Berardino Cocchiararo provided tissues for generating the positive controls. We thank Susan Mbedi from BeGenDiv for suggestions related to multiplexed sequencing library preparation and sequencing. MB is supported by DFG grant BA 4843/2-1.

## Appendices and Supplementary Materials

**Appendix 1**. Molecular ecology studies that report randomization in some part of the work. All relevant articles were screened in a randomly selected issue of four journals for the search term “random”. We deemed articles relevant when they used DNA or RNA methods that are sensitive to laboratory biases (microsatellite genotyping, SNP assays, metabarcoding, metagenomics, (meta)transcriptome comparisons, etc.). We also included also studies that use single genes for molecular identification of species or populations (e.g. barcoding and single-gene biogeographies) since identification may be non-randomly confounded by cross-contamination (a simple example would be cross-contamination when neighboring populations or related species are processed in batches). Randomization in data analysis refers to the use of mixed effect models, the generation of null hypothesis by random data rearrangements, etc.

|  | Mol. Ecol. | ISME J. | Soil Biol. Biochem. | Ecol. Evol. | J. Biogeogr. |
| --- | --- | --- | --- | --- | --- |
| Issue | 22 | 2 | 1 | 9 | 11 |
| Total relevant | 14 | 16 | 8 | 9 | 12 |
| Report randomization in sampling or data analysis | 12 | 4 | 4 | 2 | 6 |
| Report randomization in lab | 1 | 0 | 0 | 1 | 0 |

**Supplementary Files** (accessible through Figshare, https://figshare.com/s/32dbca0a906c7f06449b, DOI: 10.6084/m9.figshare.4579681): sample_infos.csv: the description of samples, negative and positive controls

multiplexing_indices.xlsx: PCR plate setup and nucleotide indices used for sample multiplexing

stechlin_assigned_190915.tab: OTU abundance table

Stechlin_organohalogene.csv: organohalogene pesticide concentrations in the sediments

lab-methods_OTU_anova.RData: ANOVA table of the multispecies generalized linear model (100 bootstraps)

Lab_LV_model_40000-iter.RData: ordination results with a latent variable model (40 000 iterations)

## Author contributions statement

MB conceived the ideas. MB and HPG designed the methodology and obtained the cores. MB and OM performed the molecular laboratory work. MS and RAD performed the organohalogene measurements. MB processed the sequences, analysed the data and lead the writing of the manuscript. All authors contributed critically to the drafts and gave final approval for publication.

